## Supplementary Material for "Multidimensional Characterization of Soft-Tissue Sarcomas with FUS-TFCP2 or EWSR1-TFCP2 Fusions"

Supplementary Figure 1

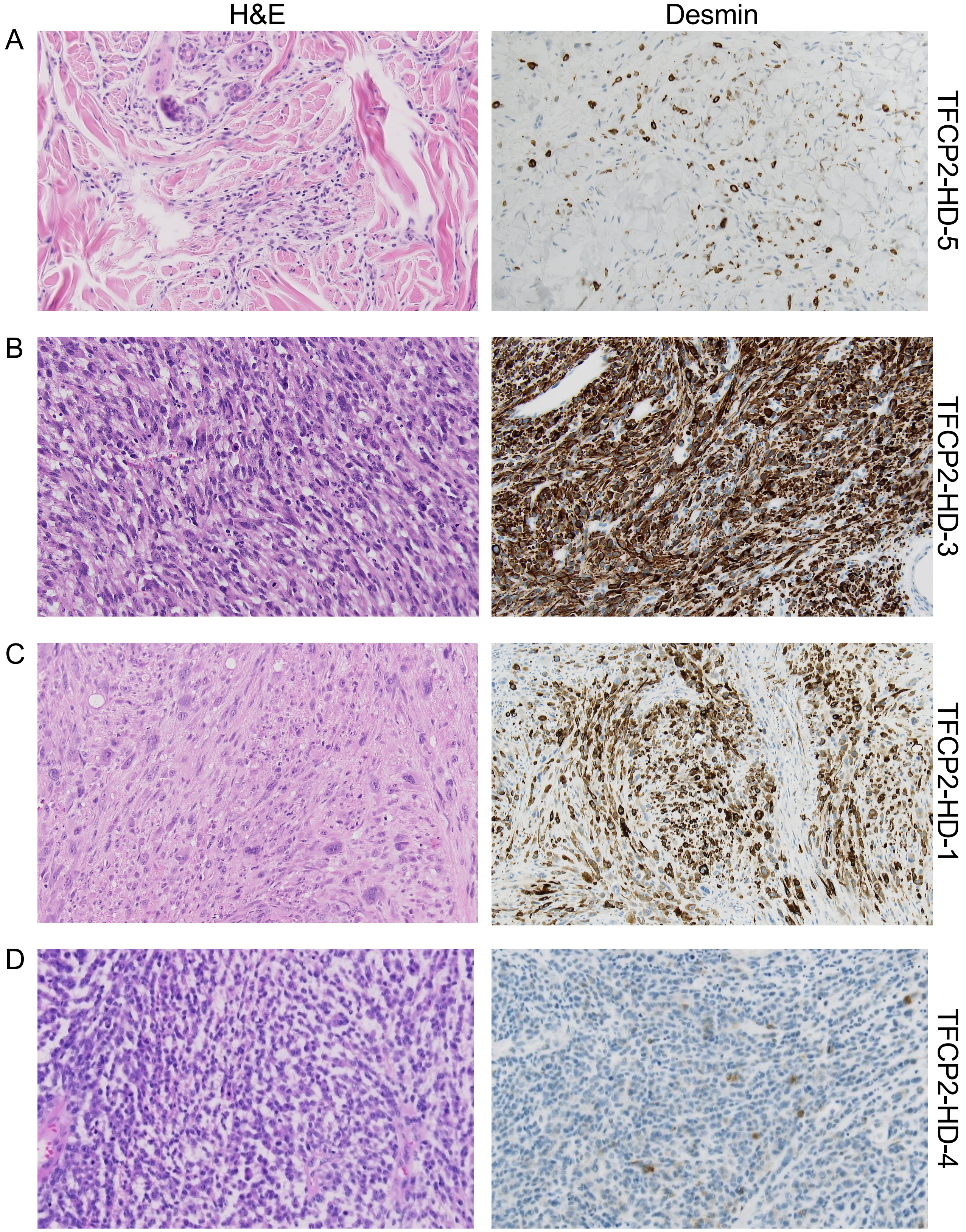


**Supplementary Figure 1. Morphologic spectrum of FUS/EWSR1-TFCP2 RMS.** FUS/EWSR1-TFCP2 RMS comprise **(A)** lesions displaying loose dermal and subcutaneous infiltrates of delicate spindle cells with few intermingled epithelioid/rhabdoid cells (pattern A), **(B)** hybrid lesions consisting of dense plump spindle cells with a quantitatively heterogeneous component of merged epithelioid/rhabdoid cells (pattern B), and **(C, D)** hybrid lesions with spindle and epithelioid/rhabdoid cells including areas with considerable pleomorphism **(C)** and/or blue cell/rhabdoid morphology **(D)** (pattern C). Images were acquired with 20x magnification.

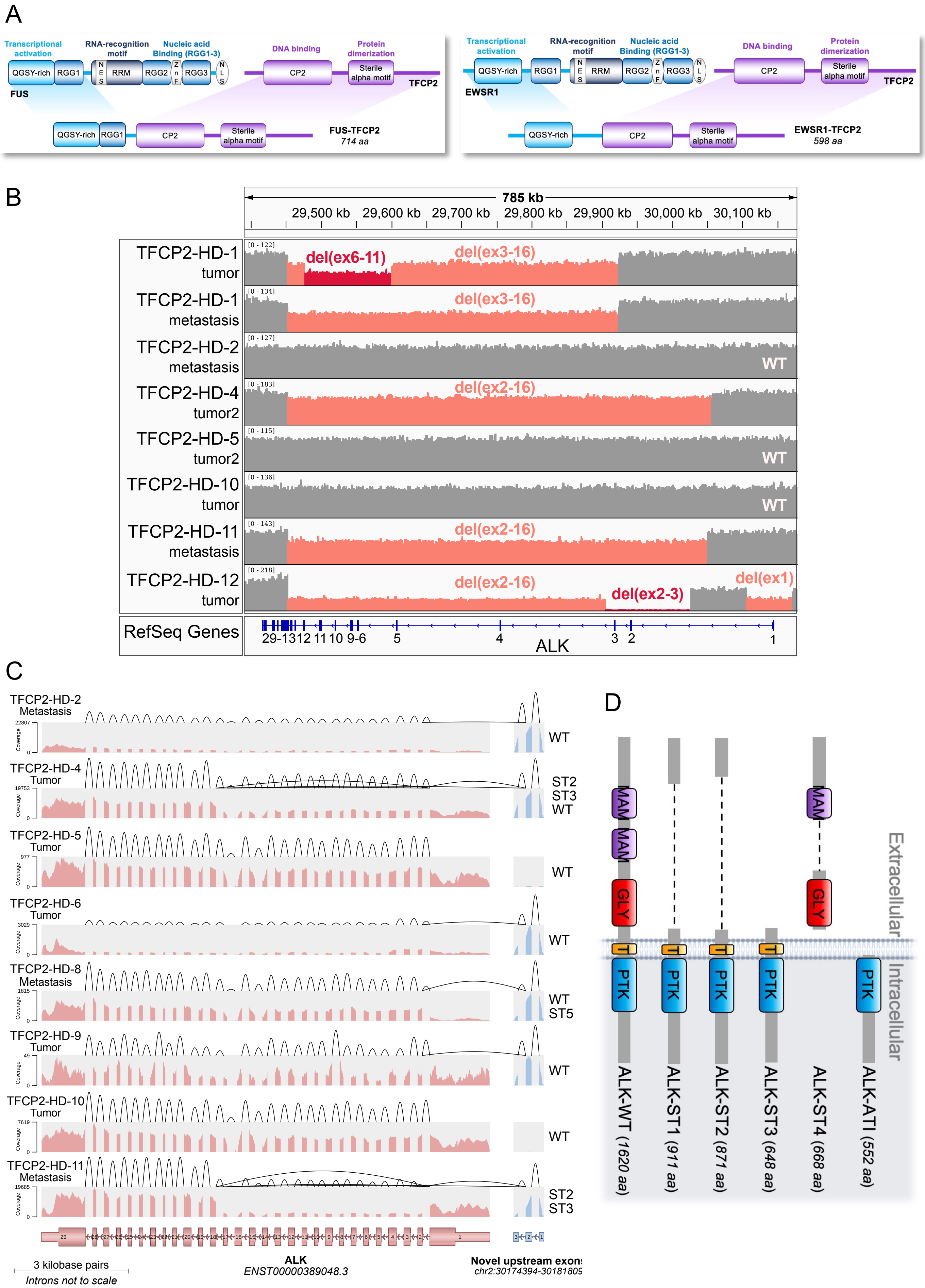
A
Transcriptional activation
RNA-recognition motif
Nucleic acid Binding (RGG1-3)
DNA binding
Protein dimerization
QGSY-rich
RGG1
RRM
RGG2
RGG3
CP2
Sterile alpha motif
FUS
TFCP2
FUS-TFCP2
714 aa
EWSR1
EWSR1-TFCP2
598 aa
B
785 kb
29,500 kb
29,600 kb
29,700 kb
29,800 kb
29,900 kb
30,000 kb
30,100 kb
TFCP2-HD-1 tumor
del(ex6-11)
del(ex3-16)
TFCP2-HD-1 metastasis
del(ex3-16)
TFCP2-HD-2 metastasis
WT
TFCP2-HD-4 tumor2
del(ex2-16)
TFCP2-HD-5 tumor2
WT
TFCP2-HD-10 tumor
WT
TFCP2-HD-11 metastasis
del(ex2-16)
TFCP2-HD-12 tumor
del(ex2-16)
del(ex2-3)
del(ex1)
RefSeq Genes
29-13 12 11 10 9-6 5 4 3 2 1
ALK
C
TFCP2-HD-2 Metastasis
WT
TFCP2-HD-4 Tumor
ST2
ST3
WT
TFCP2-HD-5 Tumor
WT
TFCP2-HD-6 Tumor
WT
TFCP2-HD-8 Metastasis
WT
ST5
TFCP2-HD-9 Tumor
WT
TFCP2-HD-10 Tumor
WT
TFCP2-HD-11 Metastasis
ST2
ST3
3 kilobase pairs
Introns not to scale
ALK
ENST00000389048.3
Novel upstream exon:
D
Extracellular
Intracellular
MAM
GLY
TM
PTK
ALK-WT (1620 aa)
ALK-ST1 (911 aa)
ALK-ST2 (871 aa)
ALK-ST3 (648 aa)
ALK-ST4 (668 aa)
ALK-ATI (552 aa)

E

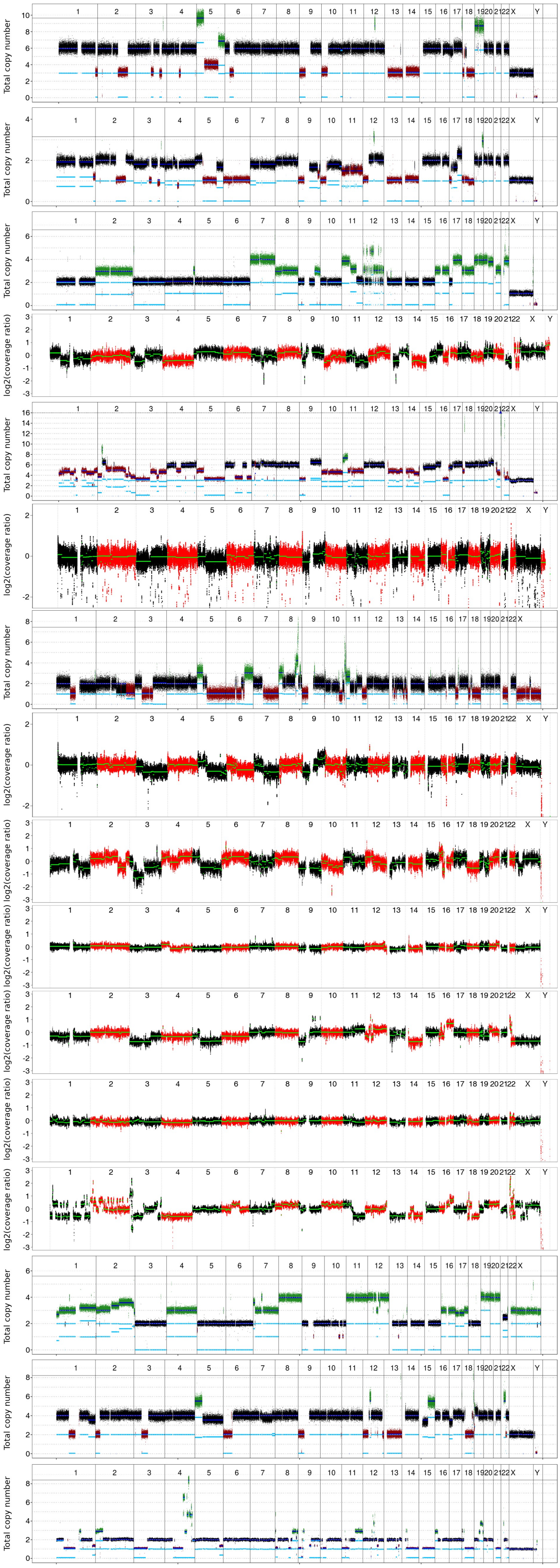

TFCP2-HD-1 Tumor
HRD-LOH: 11
**HRD-LST: 12**
HRD-TAI: 7
Score average: 10

TFCP2-HD-1 Metastasis
HRD-LOH: 10
**HRD-LST: 14**
HRD-TAI: 5
Score average: 9.7

TFCP2-HD-2 Metastasis
HRD-LOH: 4
HRD-LST: 2
HRD-TAI: 3
Score average: 3
Chromothripsis: chr12

TFCP2-HD-3 Metastasis
HRD-LOH: 9
HRD-LST: 4
HRD-TAI: 10
Score average: 7.7

TFCP2-HD-4 Tumor 2
HRD-LOH: 5
**HRD-LST: 12**
**HRD-TAI: 16**
Score average: 11

TFCP2-HD-4 Tumor 1 (early lesion)
HRD-LOH: 3
HRD-LST: 6
HRD-TAI: 3
Score average: 4

TFCP2-HD-5 Tumor 2
**HRD-LOH: 17**
**HRD-LST: 15**
**HRD-TAI: 15**
**Score average: 15.7**

TFCP2-HD-5 Tumor 1 (early lesion)
HRD-LOH: 9
HRD-LST: 3
HRD-TAI: 1
Score average: 4.3

TFCP2-HD-6 Tumor
HRD-LOH: 8
**HRD-LST: 18**
**HRD-TAI: 16**
**Score average: 14**

TFCP2-HD-7 Tumor
HRD-LOH: 0
HRD-LST: 5
HRD-TAI: 0
Score average: 1.7

TFCP2-HD-8 Metastasis
HRD-LOH: 4
HRD-LST: 7
HRD-TAI: 12
Score average: 7.7

TFCP2-HD-9 Tumor 1
HRD-LOH: 0
HRD-LST: 0
HRD-TAI: 0
Score average: 0

TFCP2-HD-9 Tumor 2
**HRD-LOH: 17**
**HRD-LST: 14**
**HRD-TAI: 17**
**Score average: 16**
Chromothripsis: chr1, chr2

TFCP2-HD-10 Tumor
HRD-LOH: 5
HRD-LST: 1
HRD-TAI: 2
Score average: 2.7

TFCP2-HD-11 Metastasis
HRD-LOH: 6
HRD-LST: 3
HRD-TAI: 4
Score average: 4.3

TFCP2-HD-12 Tumor
HRD-LOH: 15
**HRD-LST: 12**
**HRD-TAI: 15**
**Score average: 14**

F

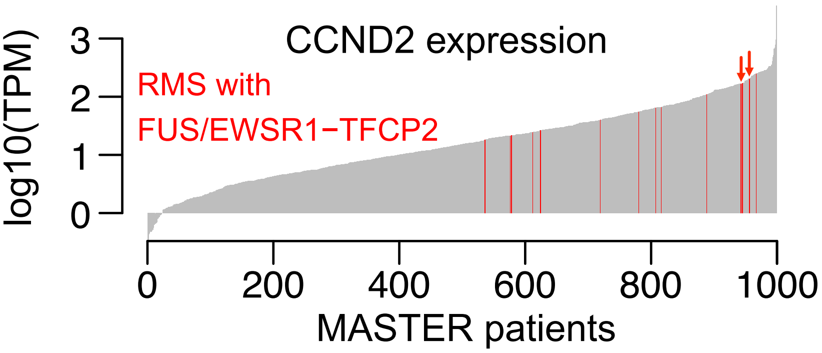


G

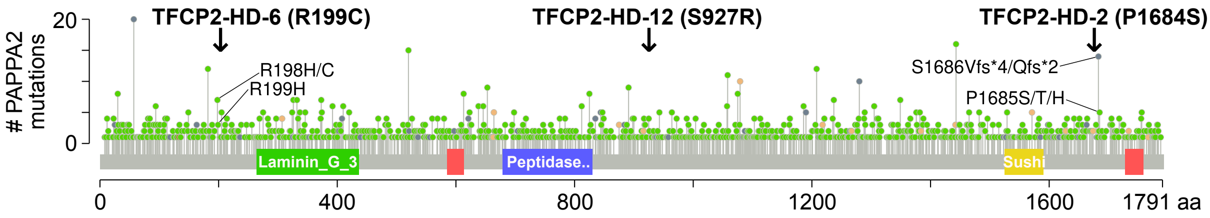


| | PolyPhen | SIFT | CADD score (PHRED-like) |
|---|---|---|---|
| PAPPA2 (R199C) | Possibly damaging | Deleterious | 22.4 |
| PAPPA2 (S927R) | Probably damaging | Deleterious | 23.8 |
| PAPPA2 (P1684S) | Benign | Tolerated | 20.3 |

H

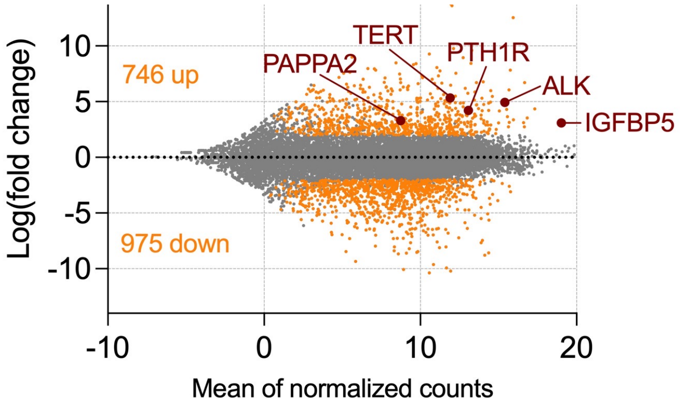


I

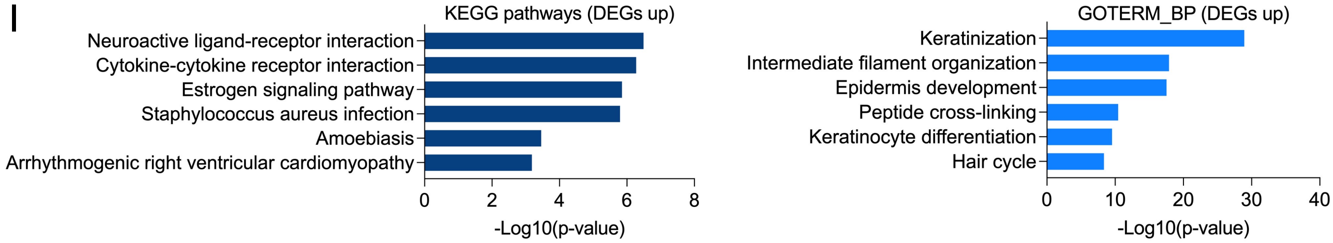


J

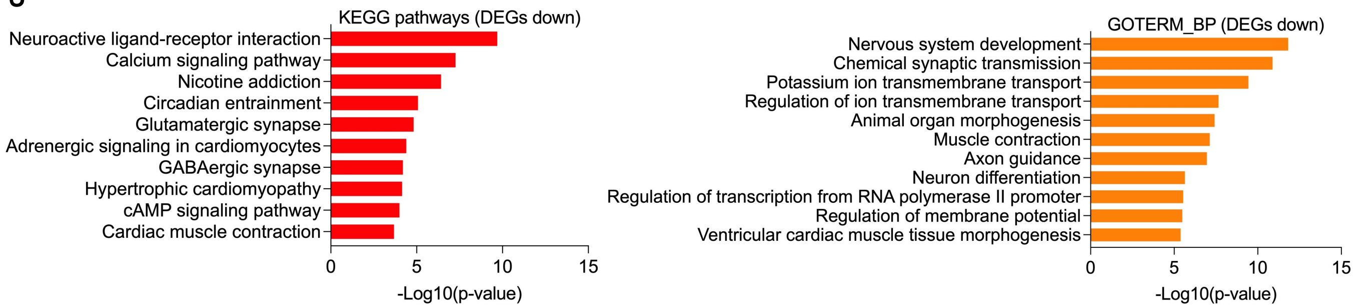

K

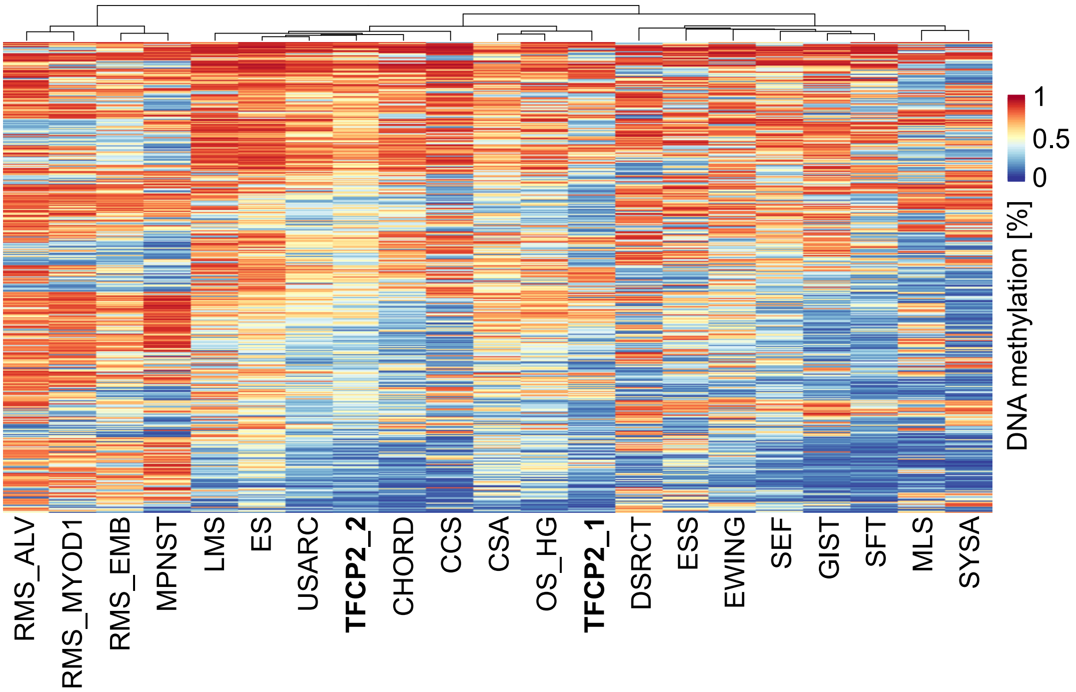


**Supplementary Figure 2. Molecular characteristics of FUS/EWSR1-TFCP2 RMS. (A)** Protein domains of FUS, EWSR1, TFCP2, FUS-TFCP2, and EWSR1-TFCP2. QGSY, Gln-Gly-Ser-Tyr-rich region; RGG, Arg-Gly-Gly-rich motif; NES, nuclear export signal; NLS, nuclear localization signal; RRM, RNA recognition motif; Znf, zinc finger motif; CP2, conserved region in CP2 transcription factor family. **(B)** ALK gene coverage plots of patient samples for which WGS data were available**.** Patient TFCP2-HD-7 carrying a del(ex2-17) is not shown because only WES data were available. Del, deletion; ex, exon; WT, wildtype. **(C)** RNA-seq data for ALK in FUS/EWSR1-TFCP2 RMS cases. The expression of ALK exons is indicated by the depth of coverage (pink, regular exons; blue, novel exons), and the expression of splice junctions is reflected by the height of the arcs connecting the exons. **(D)** Schematic of ALK-WT and the four ALK variants found in RMS. ALK-ATI has been identified in melanoma (Wiesner et al. 2015). PTK, protein kinase domain; T, transmembrane domain; MAM, meprin, A-5 protein, and receptor protein-tyrosine phosphatase mu domain; GLY, glycine-rich domain. **(E)** Copy number profiles of FUS/EWSR1-TFCP2 RMS cases. WES samples are shown in red and black colors alternating by chromosome, and detected segments are marked by green lines. WGS samples are colored by aberration type (green, gain, red, loss, black, copy number-neutral relative to base ploidy). Light blue lines show the total copy number of each allele and dark blue lines the sum of the copies of both alleles. Scores quantifying the degree of genomic rearrangement associated with HRD are indicated (LOH, loss of heterozygosity; LST, large-scale state transition; TAI, telomeric allelic imbalance). Scores in bold exceed the thresholds described in the manuscript indicating HRD. **(F)** CCND2 mRNA expression of tumors from all patients enrolled in the MASTER program until November 21, 2018, and all FUS/EWSR1-TFCP2 RMS cases (shown in red). Arrows indicate the two samples with CCND2 genomic gain. TPM, transcripts per million. **(G)** Frequency of PAPPA2 mutations in 68,088 samples from 64,959 patients in 205 curated, non-redundant studies registered in cBioPortal (cbioportal.org) together with the three mutations identified this study. PAPPA2 R199C and S927R were predicted to be deleterious by three algorithms (PolyPhen, SIFT, CADD); PAPPA2 P1684S, although near a mutational hotspot, was predicted to have no effect on protein function. **(H)** MA plot showing genes differentially expressed between FUS/EWSR1-TFCP2 RMS and other RMS cases called by DESeq2 (adjusted p-value >0.05; –log2(fold change) >2 and <–2). All differentially expressed genes with adjusted p-values >0.05 and –log2(fold change) >1 and <–1 are listed in Suppl. Table 2. **(I, J)** Top significantly enriched KEGG pathways and GOTERM_Biological Processes in up- (I) and downregulated (J) differentially expressed genes shown in (H). **(K)** Hierarchical clustering of average DNA methylation across the 6,000 most variable CpGs in TFCP2-rearranged samples (n=11; cluster TFCP2_1 and TFCP2_2) and 19 other sarcoma types. RMS_ALV, alveolar RMS; RMS_MYOD1, MYOD1-mutant spindle cell/sclerosing RMS; RMS_EMB, embryonal RMS; MPNST, malignant peripheral nerve sheath tumor; WDLS_DDLS, well-differentiated and dedifferentiated liposarcoma; SFT, solitary fibrous tumor; CCS, clear cell sarcoma; ES, epithelioid sarcoma; USARC, undifferentiated sarcoma; CHORD, chordoma; ASPS, alveolar soft part sarcoma; SEF, sclerosing epithelioid sarcoma; GIST, gastrointestinal stromal tumor; AS, angiosarcoma; CSA, chondrosarcoma; OS_HG, osteosarcoma high-grade; DSRCT, desmoplastic small round cell tumor; ESS, endometrial stromal sarcoma; EWING, Ewing sarcoma; MLS, myxoid liposarcoma; SYSA, synovial sarcoma.

Supplementary Figure 3

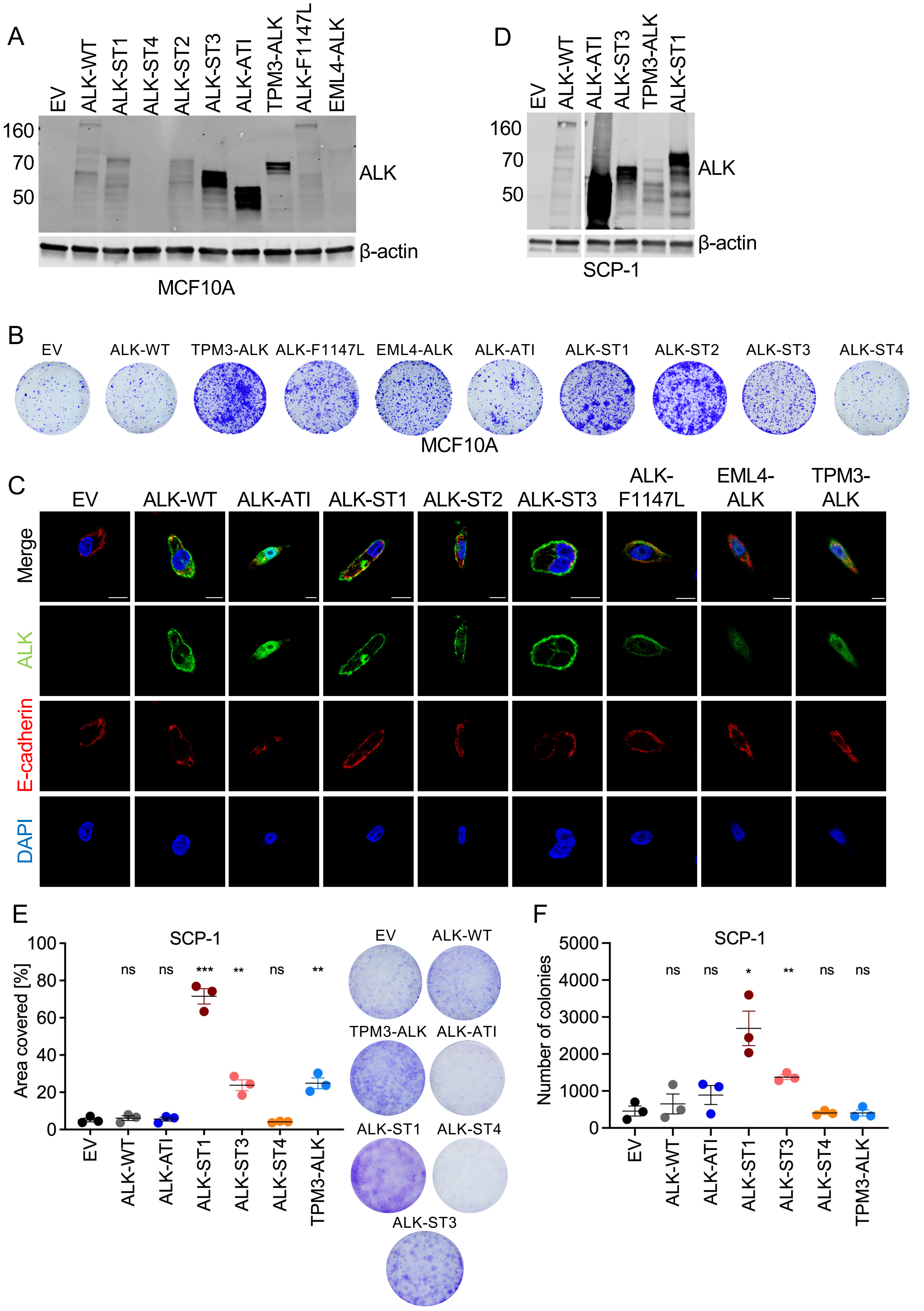

G

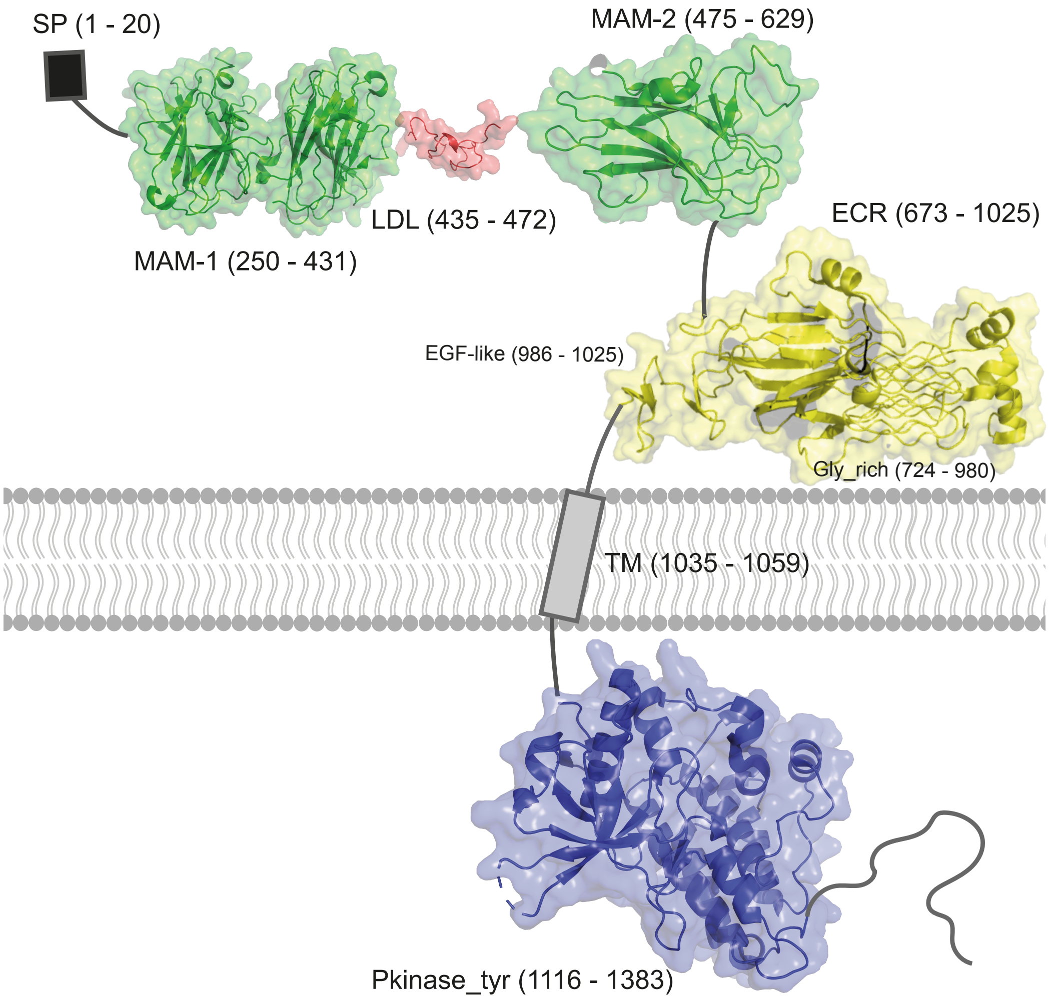


**Supplementary Figure 3. Characterization of ALK variants. (A)** Western blot of MCF10A cells stably expressing EV or ALK variants used for experiments shown in Figure 3. ALK-ST4 cannot be detected with the antibody used as it binds to the intracellular part of ALK, but expression was confirmed by qRT-PCR. **(B)** Representative images of colony formation assays with MCF10A cells stably transduced with ALK variants or EV shown in Figure 3A. **(C)** Immunofluorescence with an anti-ALK antibody (green) of MCF10A cells stably expressing EV or ALK variants. Nuclei and cell membranes were visualized with DAPI (blue) and anti-E-cadherin (red), respectively. Scale bar, 10 µm. **(D)** Western blot of SCP-1 stably expressing EV or ALK variants. The left and right parts are images of the same membrane with the same exposure from which a non-relevant lane has been cut out. **(E)** Colony formation of SCP-1 cells stably transduced with ALK variants or EV. **(F)** Anchorage-independent growth in soft agar of SCP-1 cells stably transduced with ALK variants or EV. **(G)** Modeled structure of the human ALK protein. Regions without available structures were predicted using HHpred. The structure of MAM-1 was based on the MAM domain of human neuropilin-1 (PDB: 5L73); the structure of MAM-2 was based on the beta subunit of human meprin-A (PDB: 4GWM); the structure of the ECR was based on the human ALK-ECR (PDB: 7MZW), and the Pkinase_tyr structure was based on the human ALK kinase domain (PDB: 3L9P). The structure of the short LDL receptor was predicted using MODELLER. Numbers in parentheses indicate the coordinates of the respective structural motif. SP, signal peptide; TM, transmembrane helix; Gly_rich, glycine-rich region; Pkinase_tyr, tyrosine protein kinase domain. Statistical significance was assessed by an unpaired t-test. *, $p<0.05$; **, $p<0.01$; ***, $p<0.001$.

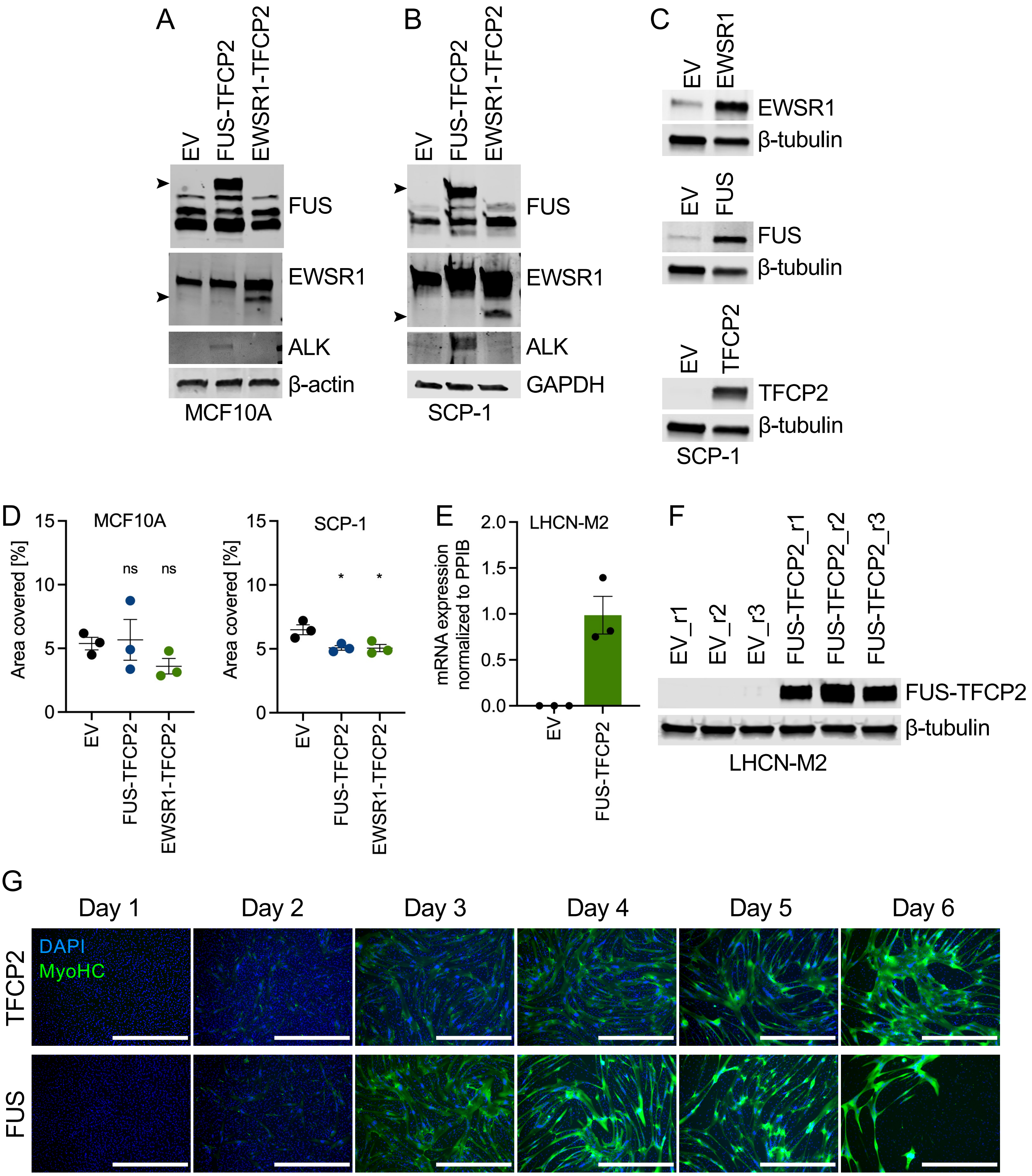


**Supplementary Figure 4. Oncogenic properties of FUS/EWSR1-TFCP2. (A-C)** Western blot of MCF10A (A) and SCP-1 (B, C) cells stably transduced with TFCP2 fusions, EV, or WT fusion partners using antibodies detecting FUS, EWSR1, and ALK. Arrowheads indicate the respective fusion proteins. **(D)** Colony formation of MCF10A and SCP-1 cells stably transduced with TFCP2 fusions or EV. **(E)** mRNA expression of FUS-TFCP2 in LHCN-M2 cells stably transduced with FUS-TFCP2 or EV. Mean ± SEM of three biological replicates. **(F)** Western blot of LHCN-M2 cells stably transduced with FUS-TFCP2 or EV in three biological replicates (r1-r3). An anti-FUS antibody was used to detect the fusion protein. **(G)** Immunofluorescence images of LHCN-M2 cells transduced with TFCP2 or FUS and cultured in differentiation medium over six days in parallel with the cells shown in Figure 4C. Green, myosin heavy chain; blue, DAPI (nuclei). Scale bar, 1 mm.

### Supplementary Figure 5

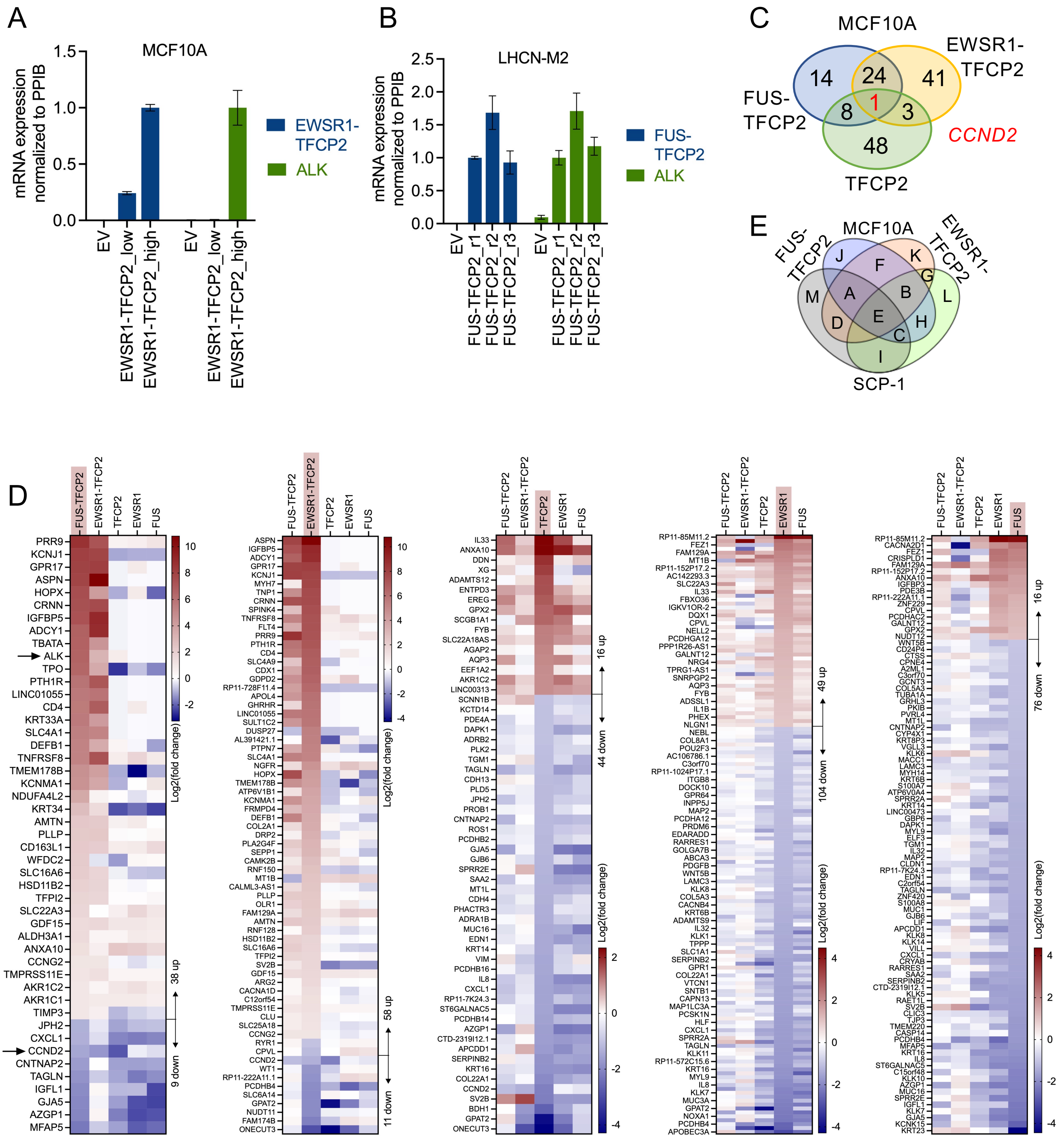

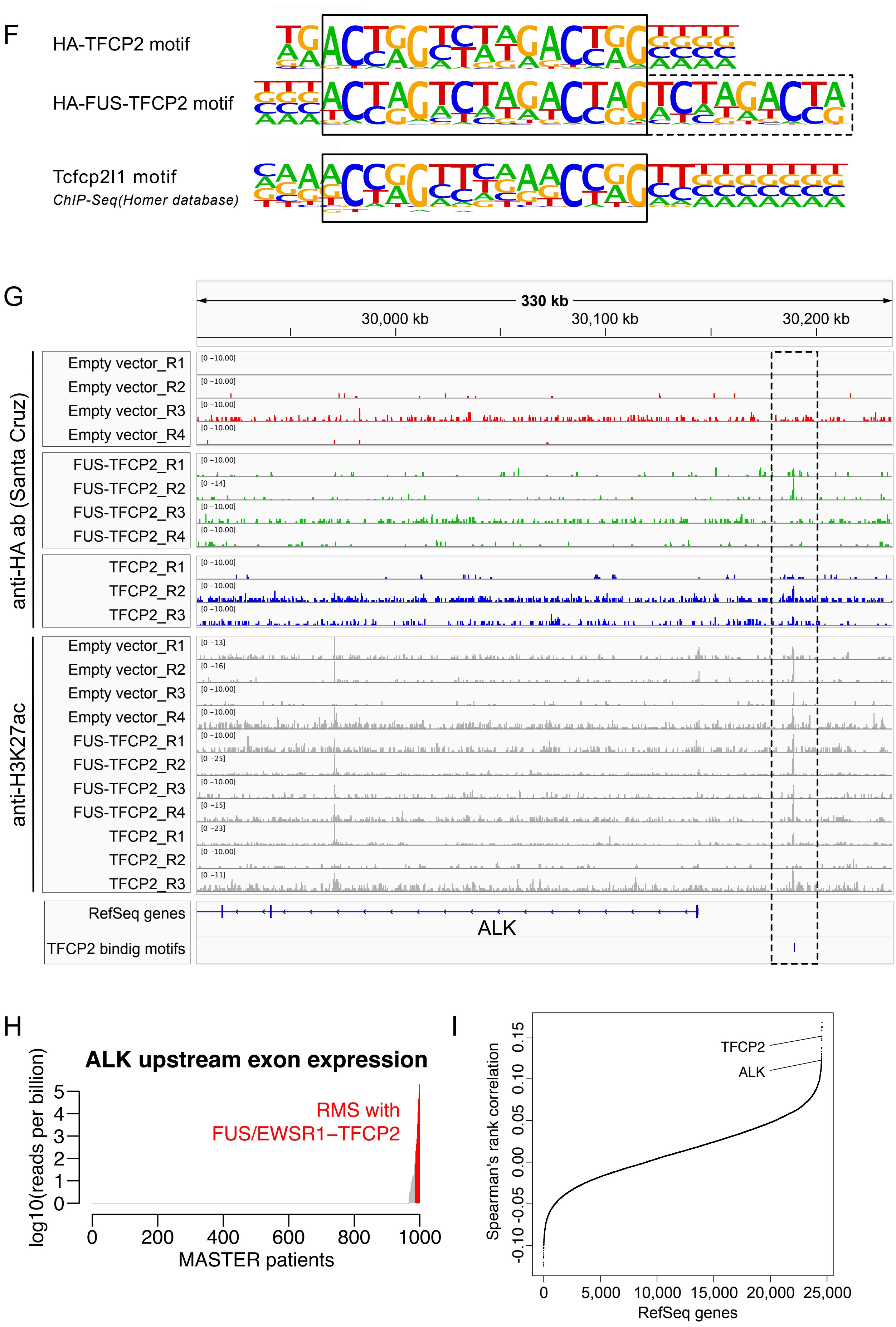
F
HA-TFCP2 motif
HA-FUS-TFCP2 motif
Tcfcp2l1 motif
ChIP-Seq(Homer database)
G
330 kb
30,000 kb
30,100 kb
30,200 kb
anti-HA ab (Santa Cruz)
Empty vector_R1
Empty vector_R2
Empty vector_R3
Empty vector_R4
FUS-TFCP2_R1
FUS-TFCP2_R2
FUS-TFCP2_R3
FUS-TFCP2_R4
TFCP2_R1
TFCP2_R2
TFCP2_R3
anti-H3K27ac
Empty vector_R1
Empty vector_R2
Empty vector_R3
Empty vector_R4
FUS-TFCP2_R1
FUS-TFCP2_R2
FUS-TFCP2_R3
FUS-TFCP2_R4
TFCP2_R1
TFCP2_R2
TFCP2_R3
RefSeq genes
TFCP2 bindig motifs
ALK
H
ALK upstream exon expression
log10(reads per billion)
RMS with FUS/EWSR1−TFCP2
MASTER patients
0
200
400
600
800
1000
I
Spearman's rank correlation
TFCP2
ALK
RefSeq genes
0
5,000
10,000
15,000
20,000
25,000

J

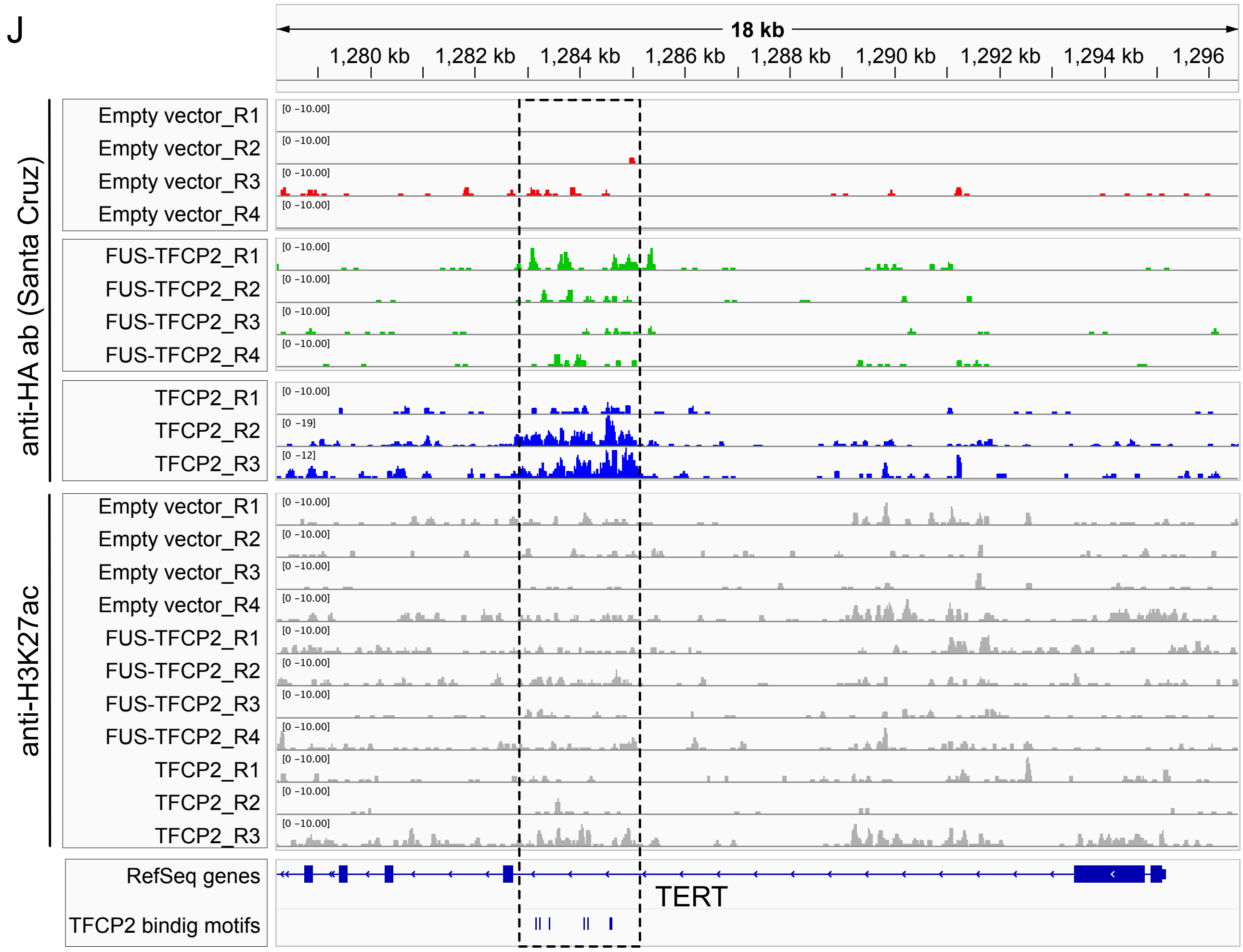


**Supplementary Figure 5. Transcriptional effects of FUS/EWSR1-TFCP2. (A)** mRNA expression of EWSR1-TFCP2 and ALK in MCF10A cells stably transduced with EV or EWSR1-TFCP2 in two biological replicates (low/high). Mean ± SEM of three technical replicates. **(B)** mRNA expression of FUS-TFCP2 and ALK in LHCN-M2 cells stably transduced with EV or FUS-TFCP2 in three biological replicates (r1-r3). Mean ± SEM of three technical replicates. **(C)** The number of genes significantly ($p<0.05$) deregulated in MCF10A cells transduced with FUS-TFCP2, EWSR1-TFCP2, or TFCP2 versus cells transduced with EV (log2(fold change) >1.0 or <–1.0) as determined by RNA-seq. **(D)** Heatmaps of genes deregulated in MCF10A cells expressing FUS-TFCP2 (first), EWSR1-TFCP2 (second), TFCP2 (third), EWSR1 (fourth), and FUS (fifth) versus cells transduced with EV. Significantly deregulated genes with log2(fold change) >1.0 or <–1.0 are shown for the cell line marked in red. **(E)** Venn diagram as in Figure 5B but with category letters indicated as they appear in Supplemental Table 2. **(F)** Binding motifs enriched in ACT-seq peaks of samples expressing HA-TFCP2 or HA-FUS-TFCP2 and comparison with the known Tcfcp2l1 motif determined by murine ChIP-seq experiments (http://homer.ucsd.edu/homer/motif/motifDatabase.html). **(G)** ACT-seq peaks detected upstream of ALK with anti-HA (Santa Cruz) and anti-H3K27ac antibodies (see Figure 5E for results with an anti-HA antibody from Cell Signaling). Enrichment of sequencing reads 45 kb upstream of ALK in cells transduced with FUS-TFCP2 or TFCP2 coincides with the transcriptional activation mark H3K27ac and a TFCP2-binding motif (dotted rectangle). **(H)** Expression of unannotated exons upstream of ALK (chr2:30174394-30181809) in tumors from all patients (gray bars) enrolled in the MASTER program until November 21, 2018, and all FUS/EWSR1-TFCP2 RMS cases (red bars) measured in units of reads per billion mapped reads. **(I)** Spearman's rank correlation of the expression of the unannotated exons upstream of ALK and all RefSeq genes computed from samples of the MASTER cohort, excluding TFCP2-rearranged tumors. TFCP2 was among the genes with the strongest correlation of all RefSeq genes. **(J)** ACT-seq peaks detected in the second intron of TERT with anti-HA (Santa Cruz) and anti-H3K27ac antibodies (see Figure 5F for results with the anti-HA antibody from Cell Signaling). Enrichment of sequencing reads near the beginning of exon 3 in cells transduced with FUS-TFCP2 or TFCP2 coincides with a cluster of TFCP2-binding motifs in this region (dotted rectangle).

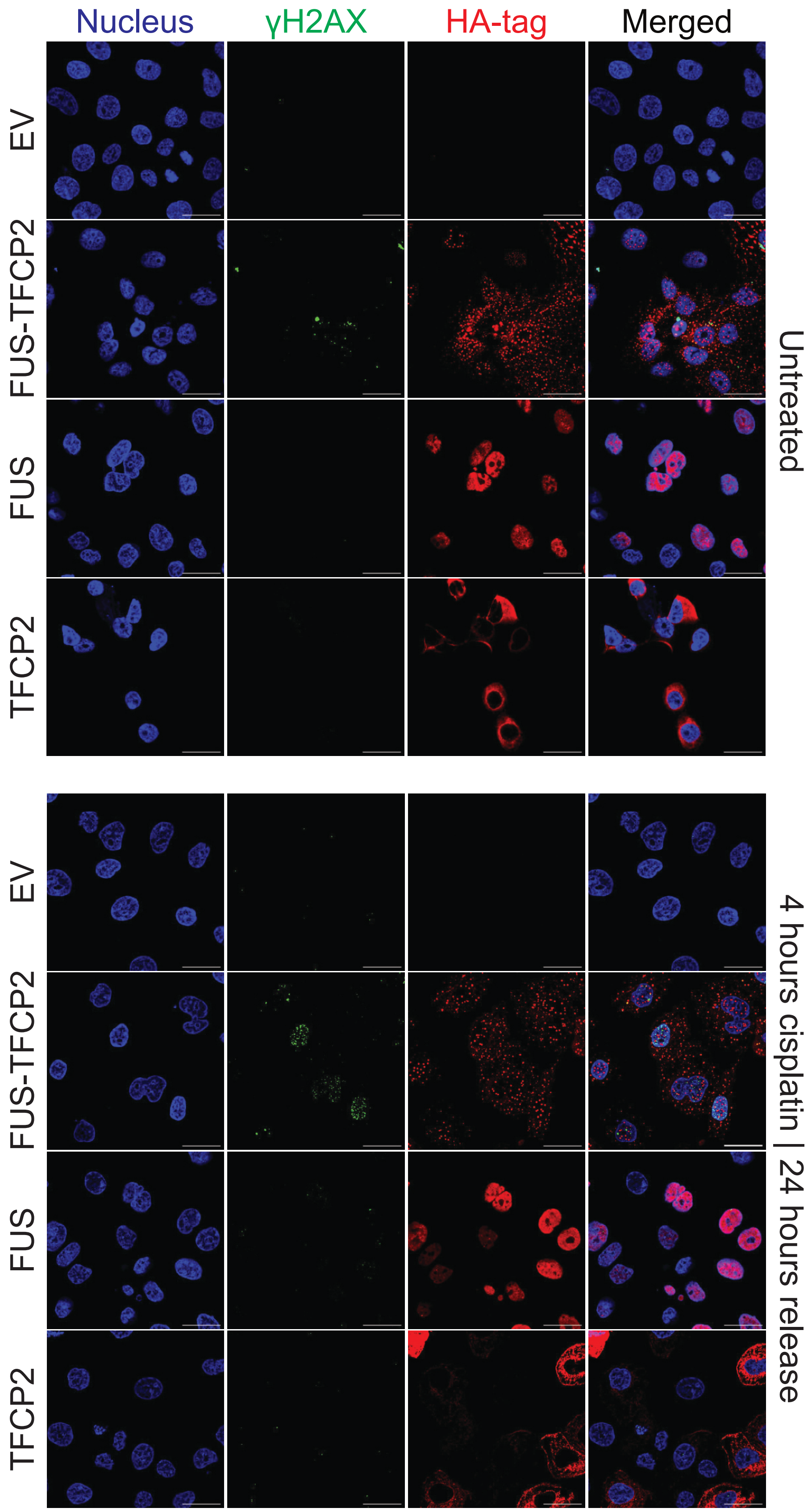


**Supplementary Figure 6. Effect of cisplatin on DNA damage repair.** Immunofluorescence images of MCF10A cells transduced with EV or HA-tagged FUS-TFCP2, TFCP2, or FUS. Cells were cultured in regular medium (untreated) or treated with 2.5 μg/ml cisplatin for four hours, followed by 24 hours in medium without drug. Details of these images are shown in Figure 6F. Scale bar, 100 μm.

### SUPPLEMENTARY TABLES PROVIDED AS EXCEL FILES

**Supplementary Table 1:** Characteristics of patients with FUS/EWSR1-TFCP2 RMS

**Supplementary Table 2:** Differentially expressed genes in FUS/EWSR-TFCP2 RMS versus other RMS subtypes

**Supplementary Table 3:** Differentially expressed genes in MCF10A and SCP-1 cells stably transduced with empty vector (EV), FUS-TFCP2, EWSR1-TFCP2, TFCP2, FUS, or EWSR1

### SUPPLEMENTARY METHODS

#### Histopathologic analysis

Immunohistochemical staining was performed with a Bench-Mark ULTRA Autostainer (VENTANA/Roche) on 3 µm tissue microarray sections. The staining procedure included heat-induced epitope retrieval pretreatment using Tris-Borate-EDTA buffer (pH 8.4; 95–100°C, 32–72 minutes), followed by incubation with primary antibodies for 16–120 minutes and signal detection employing the OptiView DAB IHC Detection Kit (VENTANA/Roche) as described previously (1). The following primary antibodies were used: Desmin (DE-R-11, prediluted, CC1 pretreatment, Cell Marque), MyoD1 (EP212, prediluted, CC1 pretreatment, Cell Marque), Myogenin (F5D, 1:200, CC1 pretreatment, Cell Marque), and ALK1 (Clone ALK1, 1:20, CC1 pretreatment, DAKO).

#### Cell culture

MCF10A (ATCC), SCP-1 (2). LHCN-M2 (Evercyte), and HEK293T cells were maintained under standard conditions and routinely tested for mycoplasma contamination. Cell line identity was verified using the Multiplex Cell Authentication Test (Multiplexion) or the Human Cell Line Authentication Service (Eurofins Genomics Germany). MCF10A TP53 knockout cells were generated by transient transfection of Cas9 and TP53 sgRNA, followed by subcloning and pooling of nine clones with confirmed gene knockout. The resulting MCF10A cells with complete p53 loss, referred to as MCF10A in the text, were used for all experiments.

#### Vectors and lentiviral transduction

The ALK, FUS, EWSR1, and TFCP2 cDNAs were obtained from the DKFZ Genomics and Proteomics Core Facility. ALK-ST1, ALK-ST2, ALK-ST4, EML4-ALK, TPM3-ALK, FUS-TFCP2, and EWSR1-TFCP2 were synthesized codon-optimized by Trenzyme. ALK-F1147L was generated by site-directed mutagenesis, and ALK-ST3 by PCR. For ACT-seq experiments, an N-terminal HA tag was added to FUS-TFCP2 and TFCP2 cDNAs by site-directed mutagenesis. cDNAs were cloned into the lentiviral expression vectors pLenti6.2/V5-

DEST (Invitrogen), pLenti CMV Puro DEST (Addgene #17452), pLEX_307 (Addgene #41392), or pLenti EF1a/TO Neo DEST (generated from pLenti CMV/TO Neo DEST [Addgene #17292] by replacing the CMV promoter with an EF1a promoter) using Gateway Technology (Invitrogen). Generation of lentiviral particles using pMD2.G (Addgene #12259) and psPAX2 (Addgene #12260) packaging plasmids in HEK293T cells and transduction of cells were performed as previously described (2). Briefly, for lentiviral transduction, MCF10A, SCP-1, and LHCN-M2 cells were seeded the day before infection in six-well plates at a density of $1.5x10^5$, $2x10^5$, and $1x10^5$ per well, respectively. The medium was removed, and 50 µl concentrated viral supernatant and 2 ml growth medium containing 8 µg/ml polybrene (Merck Millipore) were added per well. The next day, the medium was replaced with regular growth medium, and 20 hours after infection, selection was started by adding the vector and a cell line-specific concentration of an antibiotic as follows:

| Cell line | Blasticidin | Puromycin | Neomycin |
|---|---|---|---|
| LHCN-M2 | 15 µg/ml | 1.5 µg/ml | Resistant |
| MCF10A | 10 µg/ml | 3 µg/ml | 500 µg/ml |
| SCP-1 | 650 µg/ml | 2 µg/ml | 750 µg/ml |

**Quantitative RT-PCR**

Primer sequences used for quantitative RT-PCR experiments were as follows:

| Gene | Forward primer (5‘ to 3‘) | Reverse primer (5‘ to 3‘) |
|---|---|---|
| PPIB | GAGGAAAGAGCATCTACGGTG | GCTTCTCCACCTGCATCTTG |
| ALK-WT | CTGTTCAGTTGGTGGATTCGC | AAGGAGCTATGACCAGTCCC |
| ALK-ATI | GGGGGAGACCTCAAGTCCTTC | CAGCAATGTCTCGGTGGATGAA |
| ALK-ST1 | CAGAGCCAGCCAACCTAGAC | GCTTGATGTTCACTTCGCCG |
| ALK-ST2 | GGGACATCTACAGAGCCAGC | AGGATAGGGCATGTAGCCCA |
| ALK-ST3 | TTGAATACTGCACCCAGGACC | TCCCGTTTTGCCTGTTGAGA |
| ALK-ST4 | AGTACATCAGCAGCGGCAAT | TTGCTGTTCTGGTAGGCGTT |
| EML4-ALK | GCGTGATGCTGATCTGGTCT | CAGTTCCATCTGCATGGCCT |
| TPM3-ALK | GCTGAGACAAGAGCCGAGTT | CGGGGCTCTGAAGTTCCATT |
| FUS-TFCP2 | GATCTAGCGGCGGTTACGAG | GTTCTCGTTGTCAGGAGGCA |
| EWSR1-TFCP2 | AGATCCGGATGCTGGACAAC | ATGATGCCCACGCTCATAGG |
| FUS | ATAAATTTGGTGGCCCTCGG | ATCATGGGCTGTCCCGTTTT |

| | | |
|---|---|---|
| EWSR1 | AGCCTCCCACTGGTTATACT | GATAAGCAGGCTGAGTGCCA |
| IGFBP5 | GCAAGTCAAGATCGAGAGAGAC | CTCCCCCGACAAACTTGGAC |
| MYOD | AGCACTACAGCGGCGACT | GCGACTCAGAAGGCACGTC |
| MYOG | AGCCAGGGGTGCCCAG | GTCAGCCGTGAGCAGATGAT |
| TFCP2 | TGGCCGACGAAGTGATTGAA | TGCAAGGACATCACTCATGCT |

**Immunoblotting**

The following primary antibodies were used for immunoblotting:

| Name | Species | Dilution | Company | Article # |
|---|---|---|---|---|
| ALK (D5F3) XP | Rabbit | 1:2,000 in 5% milk/TBST | Cell Signaling | 3633 |
| Anti-β-Actin (AC-15) | Mouse | 1:5,000 in 5% milk/TBST | Sigma-Aldrich | A1978 |
| β-Tubulin | Rabbit | 1:1,000 in 5% BSA/TBST | Cell Signaling | 2146 |
| EWS (G-5) | Mouse | 1:100 in 5% milk/TBST | Santa Cruz | sc-28327 |
| Anti-FUS | Rabbit | 1:500 in 5% milk/TBST | Sigma-Aldrich | SAB2108528 |
| GAPDH (D16H11) | Rabbit | 1:1,000 in 5% milk/TBST | Cell Signaling | 5174 |
| HA-Tag (F-7) | Mouse | 1:200 in 5% milk/TBST | Santa Cruz | sc-7392 |
| HA-Tag (C29F4) | Rabbit | 1:1,000 in 5% milk/TBST | Cell Signaling | 3724 |
| HSP90 a/b (F-8) | Mouse | 1:200 in 5% milk/TBST | Santa Cruz | sc-13119 |
| IGFBP5 | Rabbit | 1:1,000 in 5% BSA/TBST | Cell Signaling | 10941 |
| TFCP2 (D1S3V) | Rabbit | 1:1,000 in 5% BSA /TBST | Cell Signaling | 80784 |

**Colony formation assay**

MCF10A and SCP-1 cells were seeded in six-well-plates at a density of 5,000 per well in EGF-depleted medium or 4,000 per well in normal growth medium, respectively. After eight days, cells were fixed with 100% methanol and stained with crystal violet solution (2.5% crystal violet, 20% methanol in distilled water). Plates were air-dried overnight, scanned with an Epson Perfection V850 scanner, and the area covered by cells was quantified with ImageJ and the ColonyArea macro by Guzmán et al. (3).

**Anchorage-independent growth assay**

20,000 stably transduced MCF10A or SCP-1 cells were suspended in a top layer of RPMI-1640 containing 10% FBS and 0.35% noble agar (Sigma-Aldrich) and plated on a solidified bottom layer of RPMI-1640 containing 10% FBS and 0.5% noble agar in six-well plates. The next day, 500 µl of the respective regular growth medium were added to each well and replaced every two to three days. After six weeks, colonies were stained with crystal violet solution (0.005% crystal violet, 20% methanol in distilled water), scanned with a Lionheart FX automated microscope (Bio-Tek), and counted using Gen5 software (Bio-Tek).

**Mouse experiments**

MCF10A cells expressing EV, ALK variants, or FUS-TFCP2 were harvested, washed, and resuspended in DPBS. $0.5x10^6$ MCF10A cells in 100 µl DPBS mixed 1:1 with growth factor-reduced matrigel (Corning) were injected into the flanks of isoflurane-anesthetized NOD/SCIDIL2rg$^{null}$ mice. Each cell line was applied to both flanks of three mice. After injection, animals were monitored closely, and tumor size was measured with a caliper. After reaching the maximum allowed tumor size of 1.5 cm, mice were sacrificed and tumors fixed for histopathology.

**Immunofluorescence and nuclear fusion index**

To determine the cellular localization of the different ALK variants, 50,000 stably transduced MCF10A cells were seeded on poly-D-lysine-coated coverslips in a 24-well-plate, cultured overnight, washed, fixed with 4% paraformaldehyde (PFA) in DPBS for 15 minutes at room temperature (RT), rinsed again, and blocked/permeabilized with 5% BSA and 0.1% Triton X-100 in DPBS for one hour at RT. ALK (Cell Signaling Technology, D5F3, #3633) and E-cadherin (Invitrogen, #13-1700) antibodies were diluted 1:500 in blocking buffer, added to each well, and incubated overnight at 4°C. Following three washes with 0.05% Tween-20 in DBPS, cells were incubated with goat anti-rabbit IgG Alexa Fluor Plus 488 (Invitrogen, A32731) or goat anti-mouse IgG Alexa Fluor 633 (Invitrogen, A-21052) as secondary antibodies at RT for one hour. Unbound antibody was removed by washing three times with 0.05% Tween-20 in DBPS. During the second wash, DAPI (BD Biosciences, #564907, 1:5,000) was added for counterstaining of nuclei. Coverslips with attached cells were mounted on a microscopy slide using ProLong Diamond Antifade Mountant (Thermo Fisher) and dried for 24 hours. Images were acquired with a TCS SP8 confocal microscope (Leica) and processed with ImageJ/Fiji.

To determine the nuclear fusion index, LHCN-M2 cells cultured in myogenic differentiation medium were fixed with 4% PFA in DPBS and stained using an anti-MyoHC antibody (Thermo Fisher, 14-6503-82) and DAPI as described for MCF10A cells. Thereafter,

DPBS was added to each well, and cells were imaged with a Lionheart FX automated microscope. The nuclear fusion index (%) was calculated as the number of nuclei located inside myotubes divided by the total number of nuclei within each image at 20x magnification. Myotubes were defined as MyoHC-positive with fluorescence intensity clearly above the background level and with two or more nuclei. Of each well, five images at 20x magnification were taken, and nuclei were counted manually using the multi-point tool counter in ImageJ.

**$IC_{50}$ determination of patient-derived tumor cells**

Tissue specimens were stored in MACS Tissue Storage Solution (Miltenyi) on ice, minced within 24 hours of surgery, and dissociated to single cells in Medium 199 (Sigma-Aldrich) containing Collagenase IV (Life Technologies) and calcium chloride under continuous rotation for three hours at 37° C. Digested cells were filtered with a Falcon 100 µm strainer (Corning), washed with PBS, and seeded in 384-well plates at a density of 2,000 per well in Advanced DMEM/F-12 medium (Life Technologies) + 0.6% glucose + 2 mM L-glutamine + 12 µg/ml heparin + 2% B27 (without vitamin A) + 5 mM HEPES + 10 ng/mL hFGF-basic (R & D systems, 233-FB-025) + penicillin/streptomycin. After 48 hours, cells were treated with alectinib, ceritinib, or crizotinib (Hoelzel Diagnostika) in 20 concentrations ranging from 50 µM to 0.1 nM in quadruplicates. After 48 hours of incubation, cell viability was assessed with the ATPlite luminescence-based assay (PerkinElmer). GraphPad Prism version 8.4.3. was used to generate dose-response curves with a 4PL model and to calculate $IC_{50}$ values with DMSO-treated and blank wells as negative and positive controls, respectively. Compound sensitivity was determined for each drug by relating $IC_{50}$ values with corresponding maximal reachable serum levels ($C_{max}$) (5). Cells were classified as sensitive when the $IC_{50}$ value was two-fold below the $C_{max}$, as resistant when the $IC_{50}$ value was two-fold above the $C_{max}$, and as intermediate when the $IC_{50}$ value was between the two-fold $C_{max}$ thresholds.

**RNA sequencing of cell lines**

MCF10A ($1x10^6$) or SCP-1 ($5x10^6$) cells stably transduced in triplicate (MCF10A) or three to nine replicates (SCP-1) with EV, FUS-TFCP2, EWSR1-TFCP2, FUS, EWSR1, or TFCP2 were seeded in 10- or 15-cm dishes, and RNA was isolated the next day using the RNeasy Mini Plus Kit (Qiagen). Library preparation and 125-nt paired-end read sequencing on an Illumina HiSeq 4000 system were performed at the DKFZ Genomics and Proteomics Core Facility. Six samples were pooled on the same flow cell lane, and the biological replicates were distributed on different lanes. Samples were de-multiplexed and aligned to the 1000 Genomes Phase 2 assembly of the human reference genome (hs37d5) using STAR aligner version 2.5.3a (6). Duplicate reads were marked by the markdup module of sambamba version 0.6.5 (7). The expression of genes annotated in the GENCODE version 19 gene model was quantified by the featureCounts utility of the subread package version 1.5.1 (8). RNA-SeQC version 1.1.8 was used to confirm data quality (9). A batch effect observed in MCF10A cells affecting the first replicate of each condition was corrected using the removeBatchEffect function of the limma package version 3.34.6 (10). Read counts were normalized to the sample sequencing depth and transformed using the variance-stabilizing transformation method of the DESeq2 package version 1.18.1 (11). Significantly deregulated genes were defined as genes with an adjusted p-value of ≤0.05 and a log2(fold change) of ≤–1 or ≥1.

To detect an HRD-associated gene signature in MCF10A and SCP-1 samples, RNA-seq data were compared with a published HRD expression signature consisting of 230 genes (12). LOC649679 and LOC729843 were ignored because they could not be mapped to the GENCODE version 19 gene model. Expression values were batch-corrected, normalized, and transformed as described above and converted to z-scores. Spearman's rank correlation coefficient served as a measure of how well the samples' expression profiles matched the expected HRD signature.

**Antibody-guided chromatin tagmentation sequencing**

Cells ($1x10^6$) were washed twice with cold DPBS and permeabilized in 100 µl 1x cold complex formation buffer (1x CB) (13) for 10 minutes on ice. Thereafter, 75 µl of 100% glycerol were added, and cells were either stored at –80°C or used directly. ACT-seq for histone modifications was performed according to Carter et al. (13). The pA-Tn5 transposome (pA-Tn5ome) was generated by mixing pA-Tn5ase and T5ME-A+B load adaptor mix in 2x CB. To prepare pA-Tn5ome antibody complexes, 1 µl pA-Tn5ome was mixed with 0.8 µl 1x CB and 0.8 µl antibody solution. 100,000 cells were used for pA-Tn5ome-antibody complex binding and tagmentation with anti-HA antibodies targeting HA-TFCP2 or HA-FUS-TFCP2, and 50,000 cells were used for anti-H3K27ac and IgG antibody complexes. For normalization of sequencing reads between biological replicates, approximately 3,000 permeabilized nuclei from *Saccharomyces cerevisiae* were incubated with pA-Tn5ome-antibody complex targeting yeast H2B (anti-histone H2B [yeast] HIST1H2BC rabbit mAb [BosterBio, #BOS-M30930]) and spiked into each mix of cells and pA-Tn5ome-antibody complex. Tagmented DNA purification was done with the MinElute Kit (Qiagen, #28004), and elution was performed with 22 µl of EB buffer at 55°C. To generate sequencing libraries, real-time PCR was performed in a total volume of 50 µl using 25 µl NEBNext High Fidelity 2x Mix (New England Biolabs, #M0541), 0.5 µl 100x SYBR Green, 19.5 µl tagmented eluate, and 5 µl of custom Nextera Index Primer with TruSeq UDIs (replicates R1, R3, and R4) or 2.5 µl primer T5McP1n and 2.5 µl barcode primer (replicate R2) for each sample with the following program: 72°C, 5 minutes (gap repair); 98 °C, 30 seconds (initial melting); 98°C, 10 seconds; 63°C, 10 seconds; 72°C, 10 seconds (cycling). The reaction was stopped after an increase of five or more fluorescence units.

Libraries were purified using AMPure XP beads with a bead-to-DBA ratio of 1.4:1 and 12 µl elution buffer for single-phase purification (anti-HA antibodies and IgG) and with bead-to-DBA ratios of 0.5:1 and 1.4:1 for biphasic purification (anti-H3K27ac). Library quantity and fragment size were determined using the Qubit dsDNA HS assay kit (Thermo Fisher) and a TapeStation (Agilent). For replicate R1, all samples were multiplexed and sequenced on one lane; for replicate R2, six and nine samples were multiplexed and sequenced on two lanes; for replicates R3 and R4, six to eight samples were multiplexed and sequenced on four lanes. All samples were subjected to 75-nt paired-end sequencing on an Illumina NextSeq 550 system at the DKFZ Genomics and Proteomics Core Facility. Raw sequencing data were processed with the ChIP-Seq narrow peak version 1.2.1 and ATAC-Seq broad peak version 1.2.1 pipelines of the nf-core framework (14) to detect transcription factor binding sites and histone modifications, respectively. Differential peaks were called using the edgeR module of DiffBind v2.16.2 (15), correcting for antibody- and batch-specific confounding effects whenever samples clustered by antibody or batch in principal component analysis. Enrichment of binding motifs inside ACT-seq peaks was determined with HOMER version 4.11 (16) using peaks from EV controls as background and a fixed peak size of 1,000 base pairs. Visualizations of coverage peaks were generated with Integrative Genomics Viewer version 2.12.3.

**Structural predictions**

To predict structural motifs in ALK-WT (Uniprot Identifier Q9UM73), the HHpred server was used with PDB and Pfam as target databases (17). After the initial search, which yielded significant hits in the two MAM domains, the LDL receptor class A, the extracellular region (ECR), and the tyrosine kinase domain, the structure of the short LDL receptor was predicted using models constructed with manually edited/extended alignments from HHpred and with MODELLER (18). The approximate locations of the signal peptide and the transmembrane helix were determined using Phobius (19). For predicted structural regions, the corresponding PDB files were downloaded and processed using PyMOL version 2.0. To predict the domain structure of ALK variants, the HHpred server was used with Pfam as the target database. After the initial search, which yielded the approximate location of the MAM domains, the glycine-rich region of the ECR, and the kinase domain, BLAST alignments of ALK-WT and the five ALK variants were generated to determine the location of the EGF-like domain, which is part of the ECR, in the variants. For each variant, the presence of a signal peptide and a transmembrane helix was determined using Phobius.

**DNA damage analysis**

*Cell viability and apoptosis analyses*. $0.1x10^4$ cells were seeded in white 96-well plates (Corning), and cisplatin, olaparib, or both drugs were added the next day. To measure apoptosis, cells were cultured for three days, and caspase 3/7 activity was determined using Caspase-Glo 3/7 Assay (Promega). To measure cell viability, cells were cultured for six days, and viable cells were determined using CellTiter-Glo Cell Viability Assay (Promega). For both assays, luminescence was determined with an EnVision Multimode Microplate Reader (PerkinElmer).

*Detection of γH2AX by flow cytometry*. $1x10^6$ cells were seeded in 75-$cm^2$ flasks, and treatment was started the next day. Cells were either left untreated, incubated for four hours with 2.5 µg/ml cisplatin, or treated with 2.5 µg/ml cisplatin followed by cultivation in regular growth media for 24 hours to determine the effect of cisplatin release. Cells were harvested, washed once with PBS, resuspended in 100 µl PBS, fixed and permeabilized by adding drop-wise 900 µl of ice-cold 100% methanol under gentle vortexing, and stored overnight at –20°C. Fixed cells were rehydrated by washing and incubation in 1 ml cold PBS overnight at 4°C. For staining, $0.3x10^6$ cells were resuspended in 35 µl PBS containing 1% BSA and incubated with 5 µl Alexa Fluor 488 mouse anti-H2AX (pS139) antibody (BD Bioscience) for one hour at 4°C in the dark. Cells were then washed, resuspended in 200 µl PBS containing 1% BSA, and acquired with a FACSCelesta (BD Bioscience). Data were analyzed with FlowJo v10.7.1 (BD Bioscience).

*Detection of γH2AX by immunofluorescence.* $0.8x10^5$ cells were seeded on µ-Slide 8 well chamber slides (ibidi), and treatment was started the next day as described for detection by flow cytometry. The cells were washed with PBS, fixed with 4% paraformaldehyde for 15 minutes at RT, and washed three times with PBS. For staining, cells were first blocked with PBS containing 1% BSA for one hour at RT, and then incubated with 2.5 µl Alexa Fluor 488 mouse anti-H2AX (pS139) antibody (BD Bioscience) and 2 µl HA Tag Alexa Fluor 647-conjugated antibody (R&D systems) in PBS containing 1% BSA and 0.3% Triton X-100 overnight at 4°C in the dark. The next day, the supernatant was discarded and cells were incubated with 250 µl PBS containing 1 µg/ml DAPI (BD Bioscience) for 5 minutes at RT in the dark. The cells were then washed three times with PBS and stored at 4°C in PBS until imaging. Images were taken with a Leica TCS SP8 confocal microscope (Leica Microsystems). γH2AX foci quantification was performed using CellProfiler software (https://cellprofiler.org).
